# Arginine methyltransferase signalling is hyperactive in conditions of neuromuscular junction instability and muscle atrophy

**DOI:** 10.64898/2026.09.09.750242

**Authors:** Andrew I Mikhail, Sean Y Ng, Magda A Lesinski, Stephanie R Mattina, Derek W Stouth, Rozhin Raziee, Changhyun Lim, Gautham Vasam, Keir J Menzies, Stuart M Phillips, Mark A Tarnopolsky, Vladimir Ljubicic

## Abstract

**Background:** The neuromuscular junction (NMJ) is the site of communication between myofibers and α-motor neurons. Cellular and molecular mechanisms that determine, maintain, and remodel the neuromuscular synapse are poorly understood. Coactivator-associated arginine methyltransferase 1 (CARM1) post-translationally modifies target proteins by methylating arginine residues and has emerged as a key determinant of skeletal muscle biology. Methylarginine signalling is required for the maintenance and repair of the NMJ, but the direct role of CARM1 on the NMJ in health and disease remains unexplored, particularly in humans.

**Methods:** We generated *Carm1* skeletal muscle-specific knockout-out (mKO) mice to gain a basic understanding for the role of the enzyme in NMJ biology under homeostatic and denervated conditions. Additionally, we investigated CARM1 activity in severe mouse models of neuromuscular disorders (NMDs) including D2.*mdx* and *Smn^2B/-^*mice, which replicate Duchenne’s muscular dystrophy (DMD) and spinal muscular atrophy (SMA), respectively, and exhibit chronic remodelling of the NMJ. Lastly, to evaluate if methylarginine signalling is implicated during NMJ instability in human skeletal muscle, we obtained samples from healthy volunteers before and after 14 days of single leg immobilization as well as from patients with myotonic dystrophy type 1.

**Results:** Our results demonstrated that *Carm1* mRNA expression and activity are elevated (*P<0.0*5) in NMJ-enriched regions of healthy murine skeletal muscle. *Carm1* muscle-specific deletion reduced NMJ compactness (−9.3%; *P<0.05*), increased fragmentation (+33%; *P<0.05*), and disrupted the expression of synapse-specific transcripts basally and following sciatic nerve transection. In skeletal muscle from pre-clinical models of NMDs, we observed a compensatory upregulation in CARM1-dependent arginine methylation as evident by +54% and +71 increases (*P<0.05*) in asymmetric dimethylarginine (ADMA)-marked CARM1 substrates in DMD and SMA mice, respectively. Similarly, muscle CARM1 was hyperactive with increased NMJ instability during neuromuscular disuse (+22%; *P<0.05*), and disease (+30%; *P<0.05*), in humans. In a cohort of muscular dystrophy patients and healthy volunteers, elevated CARM1 signalling was negatively correlated with clinical metrics of skeletal muscle health including grip strength (*r* = −0.583; *P<0.05*) as well as positively correlated with mRNA expression of NMJ machinery such as *CHRNA1* (*r* = 0.578; *P<0.05*).

**Conclusion:** In summary, we highlight that muscle-specific CARM1 is required for maintaining NMJ morphology and transcriptional regulation. Insults to NMJ stability during muscle disuse or in myopathic conditions were associated with enhanced CARM1-mediated methylarginine signalling in mice and humans. Collectively, our findings demonstrate CARM1 as a key mediator of NMJ biology and plasticity in health and disease.

## INTRODUCTION

The neuromuscular junction (NMJ) is the site of electrochemical communication between skeletal muscle fibers and their innervating α-motor neurons [1]. Efficient bidirectional exchange of signals at the NMJ is supported by complex mechanisms that enable optimal neurotransmission [2,3]. Our understanding of these mechanisms, which develop, maintain, and remodel the neuromuscular synapse, is limited. As such, it’s important to investigate the pathways that regulate the NMJ to identify molecules that promote synaptic health, and those that contribute to disease.

Post-translational modifications, such as phosphorylation and acetylation, are essential for maintaining cellular homeostasis and response to stress [4]. Similarly, methylation of arginine residues on target proteins is an important biological reaction that regulates gene expression, signal transduction, cell fate, and other molecular processes [5]. Protein arginine methyltransferases (PRMTs) are a family of 9 enzymes that catalyze the deposition of methyl groups on arginine residues, resulting in monomethylarginine (MMA) or dimethylarginine (DMA) marks on downstream effector proteins [6]. PRMTs are subclassified into type 1 or type 2 based on their ability to add asymmetric dimethylarginine (ADMA) or symmetric dimethylarginine (SDMA) groups to target peptides, respectively, thereby altering their activity, localization, and stability [6]. Coactivator-associated arginine methyltransferase 1 (CARM1) is a ubiquitously expressed type 1 PRMT that has broad implications in cancer and cardiovascular disease [7,8]. More recently, our group, and others, demonstrated that CARM1-mediated arginine methylation is required for skeletal muscle plasticity and contractility by orchestrating the expression of atrophy gene programs, stimulating autophagy, and maintaining mitochondrial homeostasis [S1, S2, 9–12].

PRMT1 has recently been linked to maintaining NMJ health and promoting its recovery following neuronal insults [13,14]. Further, previous qualitative analysis of *Carm1* null skeletal muscle revealed an increased prevalence of structurally abnormal NMJs and reduced *in situ* force production after direct nerve stimulation, altogether suggesting an essential biological role of methylarginine modifications in NMJ health [11]. Abnormal CARM1 protein expression has been observed in neuromuscular disorders (NMDs) such as Duchenne muscular dystrophy (DMD) and spinal muscular atrophy (SMA) [15–17], while methylarginine content is upregulated in neurodegenerative disorders, including in skeletal muscle from amyotrophic lateral sclerosis (ALS) mice and patients [18]. Despite this apparent relationship between PRMT signalling and NMJ plasticity, the role of CARM1-mediated arginine methylation on the NMJ has not been investigated. Thus, the purpose of the present study was to identify the cellular and molecular mechanisms by which CARM1 impacts the maintenance and remodelling of the NMJ and to investigate the clinical relevance of the methyltransferase in conditions of neuromuscular disuse and disease.

## METHODS

### Ethical approval

All mouse experiments were approved by the local Animal Research Ethics Board in accordance with guidelines set forth by the Canadian Council on Animal Care and are listed in the investigators’ Animal Utilization Protocol no. 22-07-26. Human immobilization (no. 7935) and myotonic dystrophy type 1 (DM1; no. 7091) trials were approved by the Hamilton Integrated Research Ethics Board, conducted according to the Declaration of Helsinki and complied with the guidelines set out in the Canadian Tri-Council policy statement on ethical conduct for research involving humans. All testing and experimental procedures were done after obtaining written, informed consent from each study participant. The studies were registered on y (immobilization: NCT05369026 and DM1: NCT04187482) and the primary outcomes were previously published by Lim et al and Mikhail et al [19,20].

### Animals

*Carm1* skeletal muscle-specific knockout (mKO), D2.*mdx,* and SMA mice as well as their respective Wild-type (WT) controls were developed as previously described and are listed in the Supplemental Information [9,21–24]. For all studies involving *Carm1* mKO mice, only males were used.We have previously observed sex-specific differences in cage activity and animal behaviour in response to the manipulation of CARM1 activity [10,25], however at the time of the experiments described in the current study we were underpowered to perform sex-based analyses so we focused on male mice. This is a limitation and our future studies will aim to appropriately address sex-based differences in some of the questions in this project. Given the X-linked recessive nature of DMD, only male WT and D2.*mdx* mice were utilized while both male and female mice were included for SMA investigations. All animals were housed at McMaster University’s Central Animal Facility under a 12-hour light/dark cycle and provided food and water *ad libitum*.

### Human experiments

To investingate the effect of neuromuscular disuse on CARM1 signalling, skeletal muscle samples before and after 14 days of unilateral leg immobilization were collected from 24 young healthy men as part of a previous investigation [19]. For experiments in individuals with DM1, 13 genetically confirmed DM1 patients and 12 age-, sex- and body mass index (BMI)-matched healthy controls were subjected to skeletal muscle biopsies from the vastus lateralis using a modified Bergström needle [S3]. Additional details regarding pariticipant characteristics, experimental procedures, sample handling and others are listed in the Supplemental Information.

### Sciatic nerve transection

Animal surgeries to induce unilateral skeletal muscle denervation were performed as previously described and outlined in the Supplemental Information [9,10].

### Protein, qPCR and mRNA sequencing

Methodology for protein and mRNA isolation and quantification are listed in the Supporting Information. All primary antibodies and qPCR primers used are listed in Tables S1 and S2.

### Whole-mount immunofluorescence (IF) NMJ staining and analysis

Morphology of the pre- and post-synapse was carried out as previously described [S4] and are outlined in detail in the Supplemental Information.

### Statistical analysis

GraphPad Prism software version 10.2.3 was used for statistical analyses. A two-tailed Student’s *t*-test, and two-way ANOVA were used to examine differences. If a significant main effect and/or interaction was observed, a Tukey or Šídák’s post hoc test was performed to identify groups that are significantly different. Additional details regarding statistics for each analysis can be found in the figure captions. Statistical outliers were determined using a Grubbs’ test, and a maximum of 1 sample was removed per group. All data are expressed as mean ± SEM. Statistical significance was accepted at *p* < 0.05.

### Data availability

All relevant RNA sequencing data and correlation outputs are available in Supplementary files. Other data from this study are available from Dr. Vladimir Ljubicic upon reasonable request.

## RESULTS

### CARM1-dependent and -independent arginine methylation in synaptic and extra-synaptic regions

We first investigated the spatial expression of key PRMTs in skeletal muscle. Using publicly available data on synaptic (NMJ) and extra-synaptic (xNMJ) compartments from the tibialis anterior (TA) of 10-month-old mice [S5], we noted a significant upregulation in *Carm1* and *Prmt7* mRNA content within NMJ compared to xNMJ-enriched regions (Figure 1A). We then leveraged the specialized motor endplate organization of the diaphragm (DIA) muscle to dissect NMJ and xNMJ samples for protein analysis (Figure 1B). Specifically, we focused on CARM1-mediated signalling as we previously demonstrated that it plays an important role in overall skeletal muscle homeostasis and neurogenic-induced atrophy [9,11]. Enrichment of axonal and nerve terminal proteins, such as neurofilament (NF) and synaptophysin (Syn), as well as the post-synaptic protein rapsyn, confirmed the successful isolation of NMJ and xNMJ fractions (Figure 1C). While the abundance of CARM1 protein was similar between xNMJ and NMJ samples, direct downstream methylation targets of CARM1, including arginine 455/460-methylated polyadenylate-binding protein 1 (mPABP1^R455/460^) and R1064-methylated BRG1-associated factor 155 (mBAF155^R1064^) were greater (*P < 0.05*) in synaptically rich regions of the DIA as compared to the xNMJ (Figure 1D-1F). However, only the methylation ratio (i.e., levels of methylated relative to total) of BAF155^R1064^ tended (*P = 0.070*) to be elevated in NMJ samples (Figure 1F). Furthermore, we observed a modestly higher (*P = 0.062*) level of asymmetric dimethylarginine (ADMA)-marked CARM1 substrates (ADMA^CARM1^), and lower (*P = 0.054*) expression of MMA, in the NMJ vs. xNMJ (Figure 1D &1F). Global ADMA and symmetric dimethylarginine (SDMA) content did not differ between xNMJ and NMJ compartments (Figure 1D & 1F).

**Figure 1.**
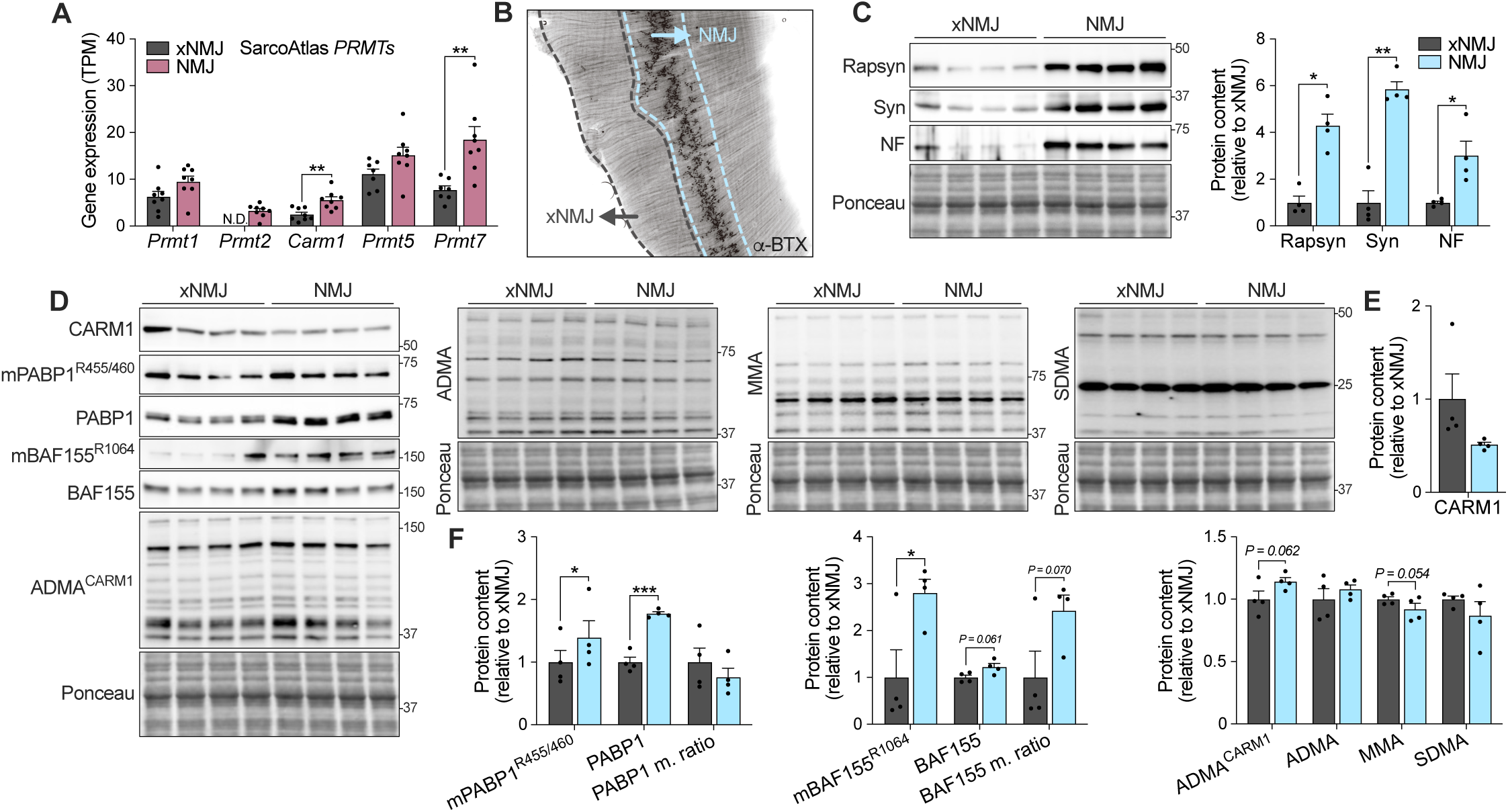
Coactivator associated arginine methyltransferase 1 (CARM1) expression and activity at the neuromuscular junction (NMJ). **(A)** Murine skeletal muscle mRNA expression of protein arginine methyltransferases (PRMTs) in extra-synaptic (xNMJ) and synaptic (NMJ) regions from the tibialis anterior (TA) of 10-month-old animals obtained from the SarcoAtlas database. n = 7-8. **(B)** Immunofluorescence microscopy inverted image of α-bungarotoxin (α-BTX) identifying the motor endplate band in mouse diaphragm (DIA) muscle. NMJ and xNMJ regions of the DIA are denoted. **(C)** Representative Western blots and graphical summary of rapsyn, synaptophysin (Syn) and neurofilament (NF) content to confirm NMJ-enrichment in DIA samples from young (18-19-week old) wild-type (WT) males. A Ponceau S stain displayed below shows sample loading and approximate molecular weights (kDa) are indicated at right of blots. n = 4. **(D)** Typical Western blots for CARM1, arginine 455/460-methylated polyadenylate-binding protein 1 (mPABP1^R455/460^), total PABP1, R1064-methylated BRG1-associated factor 155 (mBAF155^R1064^), total BAF155, asymmetric dimethylarginine (ADMA)-marked CARM1 substrates (ADMA^CARM1^), as well as total ADMA, monomethylarginine (MMA), and symmetric dimethylarginine (SDMA) in xNMJ and NMJ regions of DIA muscles. A Ponceau S stain displayed below shows sample loading and approximate molecular weights (kDa) are indicated at right of blots. **(E & F)** Summary of CARM1, methylated, total, and methylation (m.) ratio (i.e., ratio of the methylated form of the protein at the specified residue relative to the total, unmethylated protein) of PABP1 and BAF155, along with global arginine methylation markers. n = 4. Data are means ± SEM with individual data points displayed. \**P < 0.05*, \*\**P < 0.01*, \*\*\**P < 0.001* between groups. A two-tailed unpaired Student T-test was used to calculate significance.

### CARM1 is required for transcriptional and structural organization of the NMJ

Given evidence of enhanced CARM1 activity preferentially within NMJ regions, we postulated that post-synaptic CARM1 may be involved in the maintenance and remodelling of the neuromuscular synapse. Therefore, we generated *Carm1* mKO mice using a human skeletal actin (HSA)-Cre driver to deplete the protein within mature myofibers, as previously described [9,11,12,25]. *Carm1* mRNA and protein content were knocked down (*P < 0.05*) by >95% in the TA muscle of 18-19-week-old mKO male mice relative to their WT littermates (Figure 2A, S1A & S1B). Moreover, direct metrics of CARM1 activity were reduced (*P < 0.05*) by 50-70% in *Carm1* null skeletal muscle (Figure S1A & S1B). Total ADMA- and MMA-marked proteins were also significantly downregulated in mKO mice, aligning with CARM1’s role as a type 1 PRMT, whereas SDMA proteins were increased (*P < 0.05*; Figure S1A & S1B). Despite comparable gross body mass between genotypes, skeletal muscle deletion of *Carm1* resulted in a significantly smaller TA muscle mass (Figure 2B & S1C).

**Figure 2.**
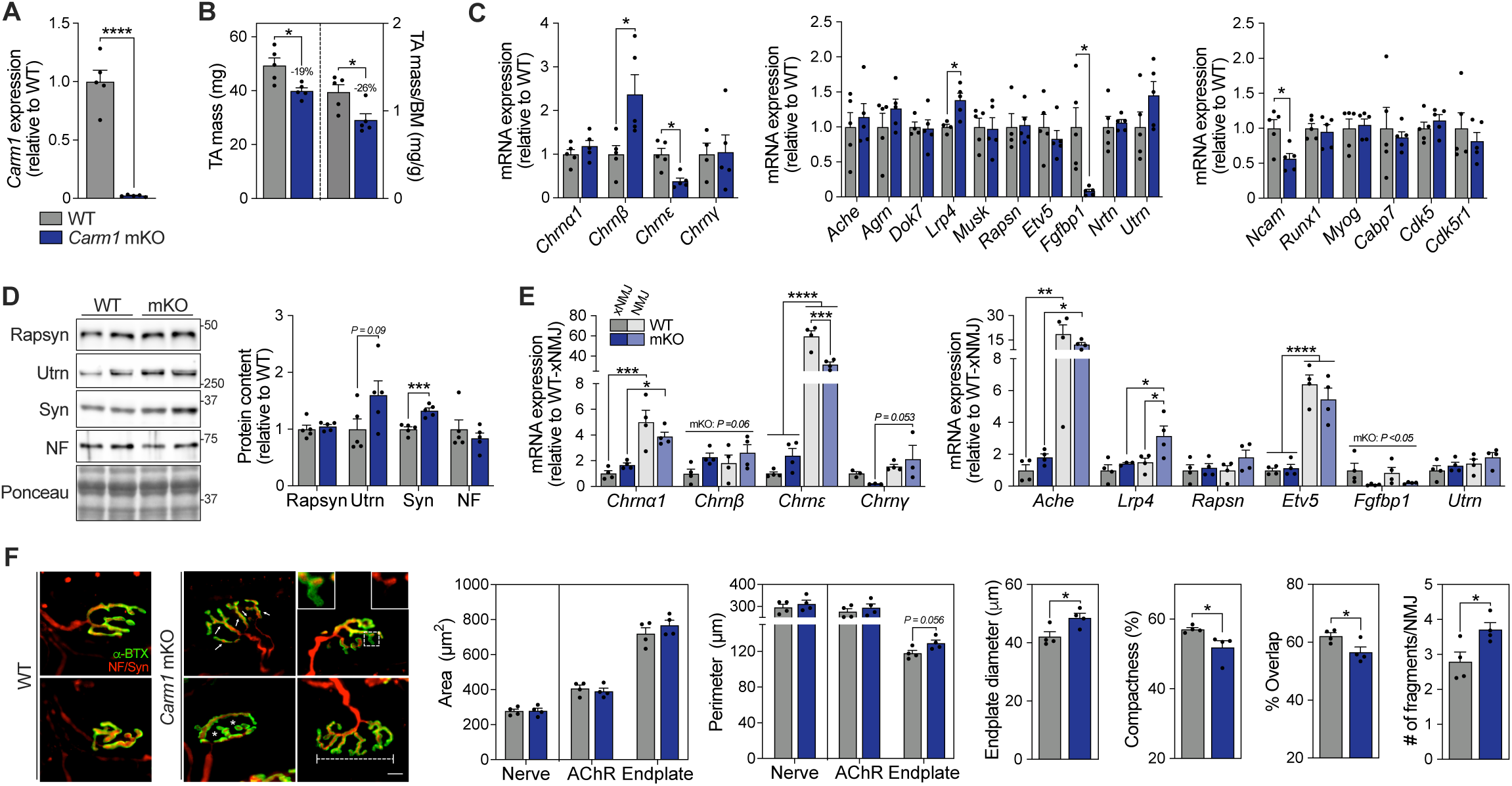
Skeletal muscle-specific deletion of *Carm1* alters NMJ gene expression and morphology. **(A)** *Carm1* transcript levels in the TA of 22-24-week old male WT and *Carm1* skeletal muscle-specific knockout (mKO) mice. n = 5. **(B)** Absolute and relative TA mass of WT and mKO mice. n = 5. **(C)** Quantification of mRNA expression of acetylcholine receptor (AChR) α1, β, ε, and γ subunits, positive regulators of the NMJ, including acetylcholine esterase (*Ache*), agrin (*Agrn*), docking protein 7 (*Dok7*), LDL receptor related protein 4 (*Lrp4*), muscle-specific kinase (*Musk*), rapsyn (*Rapsn*), ETS transcription factor 5 (*Etv5*), fibroblast growth factor binding protein 1 (*Fgfbp1*), neurturin (*Nrtn*) and *<u>Utrn</u>*, as well as denervation markers neural cell adhesion molecule 1 *(Ncam1*), runt-related transcription factor 1 (*Runx1*), myogenin (*Myog*), calcium binding protein 7 (*Cabp7*), cyclin-dependent kinase 5 (*Cdk5*), and Cdk5 regulatory subunit 1 (*Cdk5r1*) in TA muscles of WT and mKO mice. n = 5. **(D)** Typical Western blots and graphical summary of rapsyn, Utrn, Syn, and NF in TA homogenates. A Ponceau S stain displayed below shows sample loading and approximate molecular weights (kDa) are indicated at right of blots. N = 5. **(E)** Total mRNA expression of AChR subunits, and positive NMJ regulators measured in xNMJ- and NMJ-enriched DIA regions of WT and mKO mice. n = 5. **(F; Left)** Confocal microscopy images of NF and Syn (red), as well as α-BTX (green) to denote the pre- and post-synaptic compartments of the NMJ, respectively, in the epitrochleoanconeus (ETA) of WT (left) and mKO (right) mice. Scale bar = 10 µm. Arrows denote individual AChR clusters in a fragmented NMJ. Insets highlight reduced pre- and post-synaptic alignment (i.e., overlap). Asterisks indicate a greater relative area that is unoccupied by AChR (i.e., compactness). Capped line exemplifies a large endplate diameter. **(F; Right)** Metrics of pre- and post-synaptic morphology including area, perimeter, endplate diameter, compactness, overlap and fragmentation in ETA muscles. n = 4. Data are means ± SEM with individual data points displayed. \**P < 0.05*, \*\**P < 0.01*, \*\*\**P < 0.001*, *\*\*\*\*P < 0.0001* between groups. **(A-D, & F)** Two-tailed unpaired Student T-test and **(E)** two-way analysis of variance (ANOVA) followed by a Tukey or Šídák’s post hoc test, when appropriate, were used to calculate significance.

We surveyed the expression of NMJ-specific genes in whole TA muscles to broadly investigate if *Carm1* ablation alters important NMJ machinery. Transcripts encoding for the β subunit of AChR (*Chrnβ*) were ∼97% higher (*P < 0.05*) while *Chrnε* was significantly downregulated with *Carm1* deletion (Figure 2C). Other molecules including LDL receptor related protein 4 (*Lrp4*) and fibroblast growth factor binding protein 1 (*Fgfbp1*) were impacted in *Carm1* mKO mice, displaying +38% and −95% change in their expression, respectively, relative to WT counterparts (Figure 2C). Skeletal muscle CARM1 did not influence the abundance of denervation-associated genes with the exception of *Ncam*, which was blunted in mKO animals (Figure 2C). At the protein level, we observed increased utrophin (Utrn; *P = 0.09*) and Syn (*P < 0.05*) levels in TA lysates (Figure 2D). Under homeostatic conditions, the vast majority of NMJ transcripts are locally produced by a specialized population of nuclei underneath the synapse, known as fundamental, or subsynaptic, myonuclei. To understand if CARM1-mediated transcriptional deviations occur in a location-dependent manner, we examined xNMJ and NMJ regions of DIA from *Carm1* mKO and WT mice. Synaptic *Chrnε* content was ∼47% lower (*P < 0.05*) in *Carm1* mKO relative WT animals (Figure 2E). Interestingly, when taking the xNMJ regions into account, the fetal γ subunit of AChR tended to be elevated (*P = 0.053*) in NMJ regions with *Carm1* ablation. Furthermore, whole muscle upregulation of *Lrp4* in mKO mice was primarily driven by increased (*P < 0.05*) synaptic transcription (Figure 2E). Finally, we examined if altered levels of NMJ apparatus resulted in architectural remodelling of the synapse. Whole-mount epitrochleoanconeus (ETA) muscles were stained for NF, Syn and α-BTX to investigate axonal, nerve terminal, and motor endplate morphology, respectively. *Carm1* mKO resulted in larger, more fragmented NMJs with reduced compactness and overlap between pre- and post-synaptic compartments (*P < 0.05*; Figure 2F).

### CARM1 deletion alters the denervation-induced NMJ transcriptome

Next, we evaluated the role of CARM1 in the transcriptional regulation of NMJ-specific transcripts following myofiber denervation. A small portion of the sciatic nerve was excised from WT and *Carm1* mKO male mice to induce denervation (WT-Den and mKO-Den) while the contralateral limb served as a control (WT-Con and mKO-Con). TA muscles were collected 7 days post-surgery, and bulk RNA sequencing was performed (Figure 3A). Principal component analysis (PCA) revealed a robust shift in the transcriptional profile of Den TA muscles in WT and mKO mice without appreciable deviations between genotypes (Figure 3B). A total of 4,870 and 5,570 genes were differentially expressed in WT and mKO mice, respectively, in response to denervation, and ∼45% of total differentially expressed transcripts were dependent on CARM1 expression (Figure 3B; Table S3). Both WT and mKO mice demonstrated increased expression of mRNAs involved in ribosome biology and suppression of mitochondrial and oxidative phosphorylation processes (Figure 3C). Interestingly, pathways involved in synaptic translation were uniquely enriched in WT animals upon denervation, but not in the absence of *Carm1* (Figure 3C). Using a previously curated list of NMJ-specific genes from single nucleus (sn)-RNA sequencing [26], we explored the interaction between CARM1 and denervation-mediated changes in their expression. Of 472 genes, we detected 413 synaptically rich transcripts within our dataset (Figure 3D; Table S4). WT-Den and mKO-Den shared 199 mRNAs (90 up and 109 down), including canonical denervation markers (*Ncam1* and *Runx1*) and NMJ machinery (*Chrna1*, *Chrnd*, and *Ache*). Although the majority of transcripts displayed a similar directional response in both genotypes, we observed an exaggerated influence of denervation in mKO animals, as evident by significantly altered expression of transcripts involved in NMJ assembly and maintenance, such as AChR subunits (*Chrnd*, *Chrna1*, and *Chrng*) and positive regulators (*Lrp4*; Figure 3E). Additionally, ∼20% of NMJ genes were uniquely impacted by denervation in either WT or *Carm1* mKO mice, including *Lrp4* and *Lrtm1,* which have been previously investigated in skeletal muscle and NMJ biology (Figure 3F). However, the biological relevance and influence of most listed candidates on the NMJ remain largely unexplored (Figure 3F).

**Figure 3.**
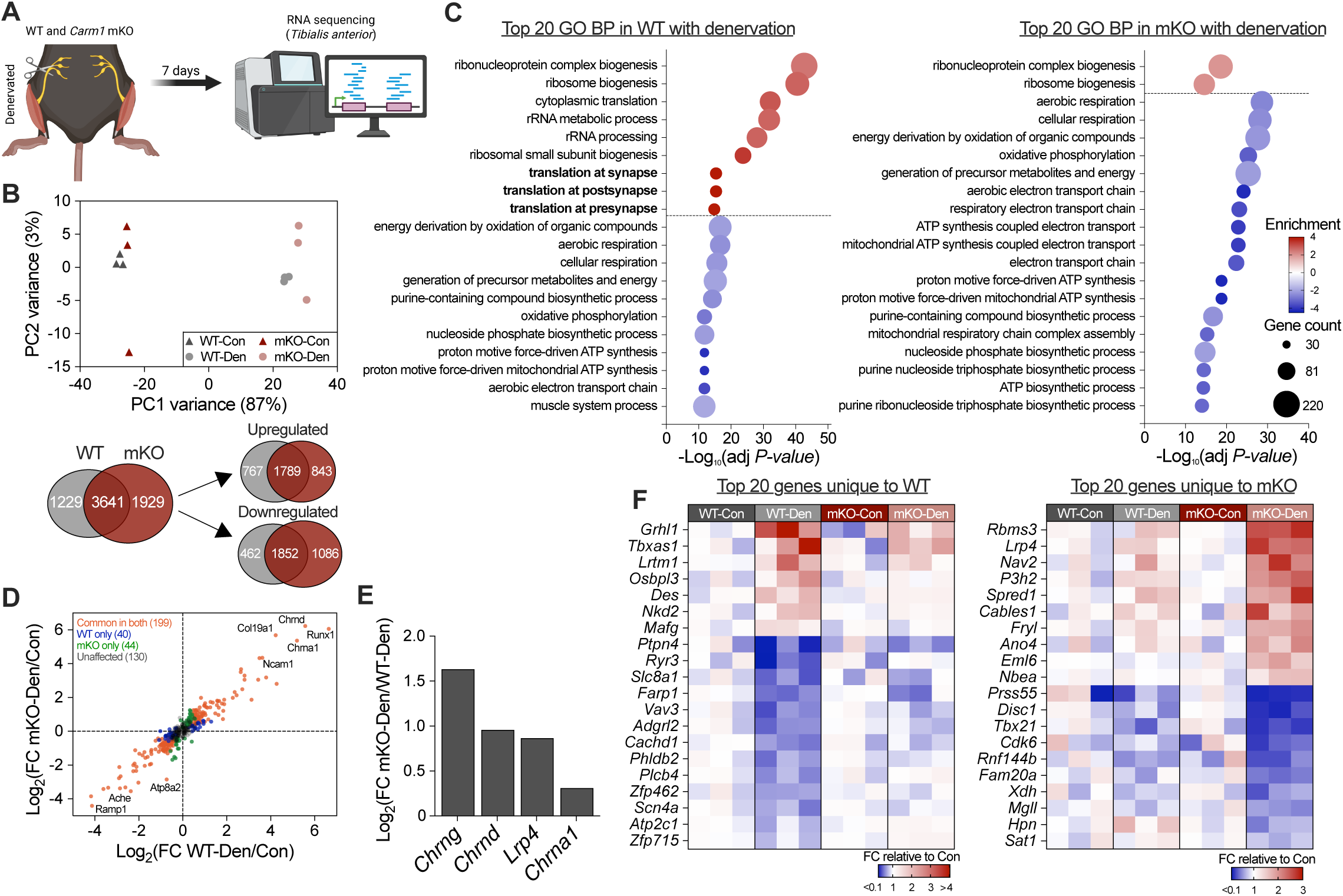
*Carm1* mKO mice exhibit a unique NMJ transcriptional profile following sciatic nerve transection. **(A)** Visual schematic of the study design. Twelve-week-old WT and *Carm1* mKO male mice were subjected to unilateral sciatic nerve transection to denervate lower limb muscles for 7 days. TA from denervated (Den) and innervated control (Con) legs were collected and subjected to RNA sequencing. (**B; Top**) Principal component analysis (PCA) to visualize the variance in bulk RNA sequencing data from WT-Con, WT-Den, mKO-Con and mKO-Den. n = 3. **(B; Bottom)** Pairwise Venn diagram summarizing differentially expressed (Log_2_(FC) > |0.5|, adj. *P*-*value* < 0.05) transcripts in WT and mKO TA muscles after denervation. **(C)** Bubble plot of the top upregulated (red) and downregulated (blue) gene ontology (GO) biological processes (BP) pathways in response to denervation in WT (left) and *Carm1* mKO (right) mice. Colour of the bubble indicates upregulated (red) or downregulated (blue) enrichment while the size highlights the number of genes represented in each pathway. (**D**) Scatterplot summarizing common and genotype-specific changes in NMJ-enriched transcripts (list derived from GSE267913) following 7 days of denervation. **(E)** Bar graph of key NMJ regulators that were significantly different between mKO-Den and WT-Den groups. **(F)** Heatmap of the top 20 uniquely altered NMJ transcripts in WT (left) and mKO (right) animals expressed as fold-change (FC) relative to their respective Con groups. A Benjamin-Hochberg (BH) multiple comparisons correction was applied and genes with an adj. *P-value* < 0.05 were considered significant.

### CARM1 hyperactivity is a common feature in skeletal muscle of NMDs

The observed NMJ structural anomalies and transcriptional remodelling induced by *Carm1* ablation led us to question whether the activity of the arginine methyltransferase is perturbed during chronic NMJ remodelling. NMDs are a group of heterogeneous conditions that arise from genetic or acquired defects within skeletal muscle, the NMJ and/or α-motor neurons [27]. DMD is among the most prevalent NMDs caused by mutations and the lack of production of functional dystrophin protein leading to fragile myofibers and continuous cycles of regeneration and degeneration along the sarcolemma including at the NMJ [28]. Using the D2.*mdx* mouse model of DMD, we confirmed the presence of abnormally large NMJs that display severe fragmentation, post-synaptic disorganization, and excessive pre-synaptic branching relative to their WT unaffected controls (Figure 4A). Concomitantly, TA muscles from D2.*mdx* mice display normal amounts of CARM1 protein but significantly greater levels of all downstream protein indicators of its activity (Figure 4B & S2A). We probed for *Fgfbp1* as a highly responsive transcriptional target of CARM1 and noted a ∼3.9-fold increased expression in dystrophic animals relative to their healthy controls (Figure 4C).

**Figure 4.**
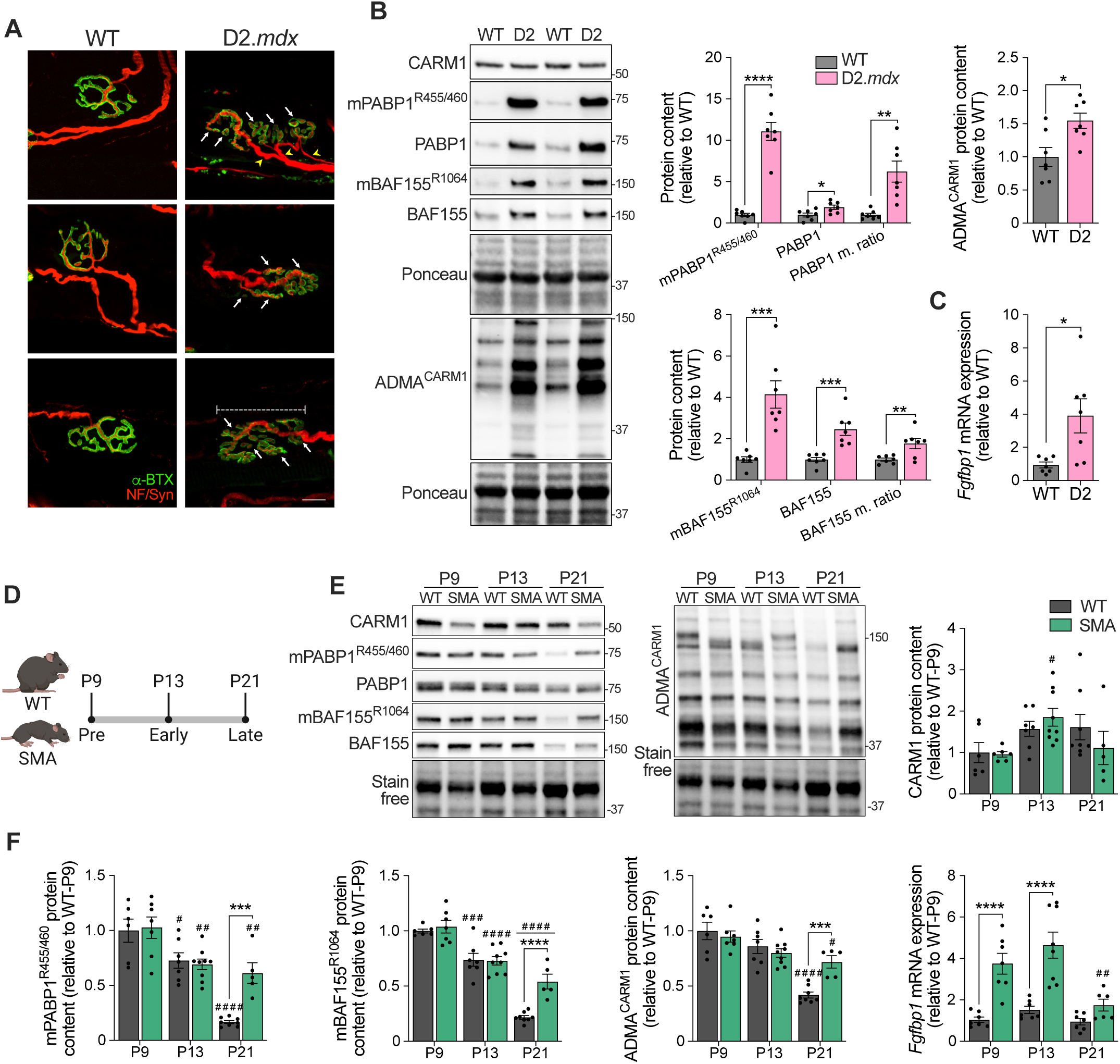
CARM1-dependent arginine methylation is upregulated in mouse models of neuromuscular disorders (NMDs). **(A)** Confocal microscopy images of NF and Syn (red) to visualize the pre-synapse and α-BTX (green) to denote post-synaptic AChRs at NMJs in ETAs of 6–8-week-old male WT and D2.*mdx* mice. White arrows denote individual AChR clusters in a fragmented NMJ. Yellow arrow heads highlight excessive axonal branching and sprouting. Capped line exemplifies a large endplate diameter. Scale bar = 10 µm. **(B)** Typical Western blots and summaries for PABP1 (methylated, total, and methylation ratio), BAF155 (methylated, total, and methylation ratio), and ADMA^CARM1^ in the TA muscles of WT and D2-*mdx* (D2) mice. Ponceau stains show sample loading and approximate molecular weights (kDa). n = 7. **(C)** *Fgfbp1* mRNA expression in the TA muscles of WT and D2-*mdx* mice. n = 7. **(D)** Schematic of the study design used to evaluate CARM1 expression and activity during spinal muscle atrophy (SMA) disease progression. Triceps (TRI) were collected from male and female WT (*Smn^2B/+^*) and SMA-like (*Smn^2B/−^*) mice at postnatal day 9 (P9), P13 and P21 to represent pre-, early- and late-symptomatic stages of the NMD, respectively. **(E)** Western blots for CARM1, mPABP1^R455/460^, PABP1, mBAF155^R1064^, BAF155 and ADMA^CARM1^ in TRI muscles of WT and SMA mice. A stain free blot displayed below indicates sample loading and approximate molecular weights (kDa) are shown at right of blots. Quantification of **(E; Right)** CARM1, **(F)** mPABP1^R455/460^, mBAF155^R1064^ and ADMA^CARM1^ protein expression in TRI muscles. n = 5-9. **(F; Left)** Total mRNA expression of *Fgfbp1* in the TRI muscles of WT and SMA mice. n = 6-8. Data are means ± SEM with individual data points displayed. \**P < 0.05*, \*\**P < 0.01*, \*\*\**P < 0.001*, *\*\*\*\*P < 0.0001* between different genotypes. ^#^*P < 0.05*, ^##^*P < 0.01*, ^###^*P < 0.001*, ^####^*P < 0.0001* between age groups within the same genotype. **(B & C)** A two-tailed unpaired Student T-test and **(E-G)** a two-way ANOVA followed by a Tukey or Šídák’s post hoc test, when appropriate, were used to calculate significance.

Next, we explored dynamic changes in CARM1 signalling in TRI muscles of a mouse model of SMA that has a well-studied disease trajectory. SMA is characterized by a deficiency in full-length survival motor neuron protein, resulting in cell-autonomous and non-autonomous defects within α-motor neurons and skeletal muscle that dismantle the NMJ and result in the loss of synaptic activity [29]. As such, skeletal muscle samples were collected from WT and SMA mice at post-natal day 9 (P9), P13 and P21 to capture pre-, early- and late-symptomatic timepoints of the disorder (Figure 4D). Of note, our group has previously demonstrated that appreciable skeletal muscle atrophy and denervation is only evident in SMA mice during late symptomatic stages (i.e., P21) [23]. We, once again, observed a disconnect between total CARM1 protein content and its enzymatic function whereby CARM1 remained stable in WT and SMA across all ages, while mPABP1^R455/460^, methylation ratio of PABP1^R455/460^, mBAF155^R1064^, and ADMA^CARM1^ levels were augmented (*P < 0.05*) by 71-260% in SMA relative to WT mice at peak disease severity (Figure 4E, 4F & S2B). Meanwhile, *Fgfbp1* was significantly higher during pre- and early-symptomatic stages of SMA (Figure 4F).

### Skeletal muscle disuse stimulates AChR mRNA expression and CARM1-dependent methylation in healthy humans

To explore the translatability of our findings, we defined the changes in CARM1 activity in response to immobilization-induced NMJ instability in young healthy volunteers. A unilateral knee brace was used to limit leg mobility for 2 weeks in male participants, and skeletal muscle biopsies from the vastus lateralis were collected pre- and post-intervention (Figure 5A) [19]. Transcripts encoding for the α1, β1, δ and γ subunits of AChR were significantly increased post-immobilization (Figure 5B). Similarly, publicly available RNA sequencing data, summarized using MetaMEx, supports upregulation (P < 0.05) of *CHRNA1*, *CHRNB*, and *CHRND* following disuse across different studies and modalities (Figure 5C) [S6]. In contrast, mRNA content of the most abundant *PRMTs* in skeletal muscle remained unaffected (Figure 5D). We attempted to measure CARM1 concentrations in protein lysates from human skeletal muscle but were unable to produce reliable data due to the presence of non-specific bands. Nevertheless, methylation targets of CARM1 were significantly enhanced in Post samples (Figure 5E), along with ADMA^CARM1^, ADMA and MMA (Figure 5F). Interestingly, SDMA marks, which are primarily catalyzed by type II PRMTs, were unresponsive to immobilization (Figure 5F).

**Figure 5.**
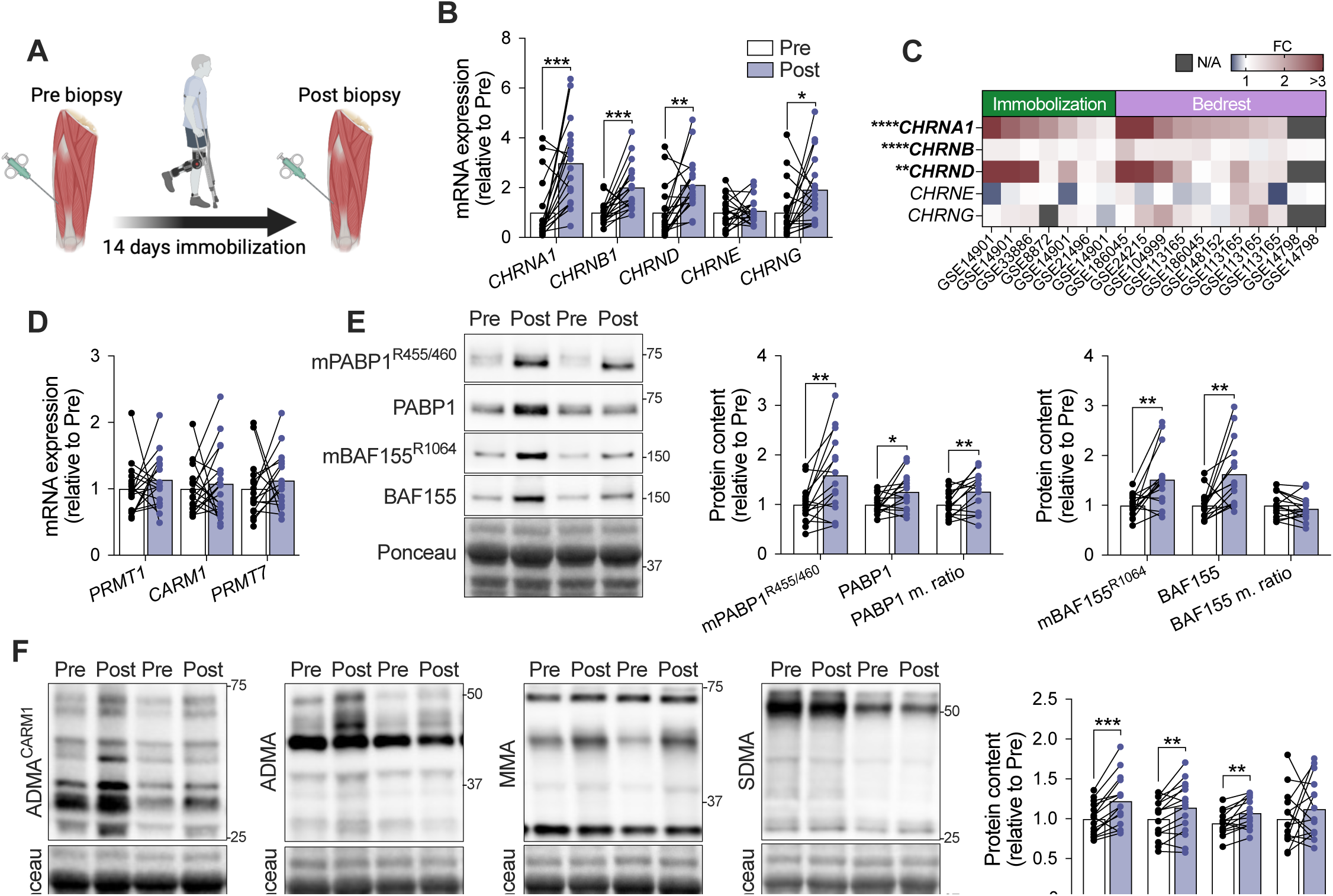
Reduced neuromuscular activity induces NMJ instability and CARM1 signalling in human skeletal muscle. **(A)** Visual schematic of the study design. Young healthy male volunteers were subjected to 14 days of single leg immobilization using a fixed knee brace. Skeletal muscle biopsies from the vastus lateralis were obtained before (Pre) and after (Post) the 2-week intervention. **(B)** Total mRNA expression of AChR subunits from Pre and Post biopsies. n = 18. **(C)** Heatmap summarizing AChR transcript expression in human skeletal muscle samples following immobilization and prolonged (i.e., 5-90 days) bedrest, an alternative model of disuse. Results were acquired from the MetaMEx platform. **(D)** mRNA abundance of *PRTM1, CARM1* and *PRMT7* in muscle biopsy samples obtained Pre and Post immobilization. n = 15-18. **(E)** Representative Western blots for mPABP1^R455/460^, PABP1, mBAF155^R1064^, and BAF155. Ponceau S stains displayed below show sample loading with approximate molecular weights (kDa). Corresponding graphical summaries for PABP1 (methylated, total and methylation ratio), and BAF155 (methylated, total and methylation ratio). **(F)** Typical Western blots and summary for ADMA^CARM1^, ADMA, MMA, and SDMA. Ponceau S stains displayed below show sample loading with approximate molecular weights (kDa). n = 14-15. Data are means ± SEM with individual data points displayed. \**P < 0.05*, \*\**P < 0.01*, \*\*\**P < 0.001*, *\*\*\*\*P < 0.0001* between groups. **(B & D-F)** A two-tailed paired Student T-test was used to calculate significance.

### Myotonic dystrophy type 1 (DM1) patient skeletal muscle is characterized by elevated NMJ transcripts and CARM1 signalling

Finally, we focused on the implications of CARM1-mediated arginine methylation on conditions of chronic neuromuscular insults in humans with NMDs. Specifically, we examined samples from patients with DM1, which is the most prevalent muscular dystrophy in adults. The pathophysiology of DM1 is driven by a trinucleotide repeat expansion mutation that alters the function and localization of RNA-binding proteins, disrupting genome-wide splicing. Emerging evidence in pre-clinical models of DM1 suggests that NMJ abnormalities are a contributing factor to its myopathic features [30–32]. We first questioned whether DM1 patients exhibit synaptic instability by confirming the presence of a denervation-like transcriptional signature in skeletal muscle from a group of genetically confirmed DM1 patients compared with unaffected age-, sex- and BMI-matched controls (Con; Figure 6A). Targeted assessment of NMJ-enriched genes from bulk RNA sequencing revealed 126 increased and 87 decreased transcripts in DM1 skeletal muscle relative to Con (Figure 6B; Table S5). Upregulated mRNAs were associated with NMJ development, synaptic assembly and ACh signalling; meanwhile downregulated genes were involved in endoplasmic reticulum stress and neuron differentiation (Figure 6B). In line with these data, DM1 skeletal muscle had greater (*P < 0.05*) levels of transcripts encoding AChR subunits, molecules that promote NMJ health, and denervation-sensitive genes (Figure 6B). In contrast, *RAPSN* expression was significantly blunted in DM1 compared to Con (Figure 6B). When we surveyed the levels of all PRMTs in muscle, *CARM1* was most affected in DM1, showing a ∼50% reduction (*P < 0.05*; Figure 6B). Additionally, we observed contrasting results showing significantly lower levels of mPABP1^R455/460^ and its methylation ratio, as well as higher expression of mBAF155^R1064^ and total BAF1555 (Figure 6C). However, ADMA^CARM1^ and non-specific ADMA marks were enhanced (*P < 0.05*) in skeletal muscle of DM1 patients (Figure 6D). In an attempt to broadly evaluate if CARM1’s function as a transcriptional co-activator is also impacted in DM1 skeletal muscle, we generated a short list of genes that were downregulated by >2-fold in the contralateral limb of mKO compared to WT mice within our RNA sequencing analysis in Figure 3, and are therefore likely to be transcriptionally regulated by CARM1. Of 20 genes identified, 15 transcripts were detected in our DM1 dataset after low count filtration, 4 of which were significantly upregulated in DM1 (i.e., CDH4, HDAC9, HS3ST5, TAS1R1) while the remaining 11 were unchanged (Figure 6E). To establish a relationship between CARM1, DM1 disease phenotype and NMJ instability, we performed a *Pearson’s* correlation on ADMA^CARM1^ as a direct marker of CARM1’s global enzymatic activity against several metrics of skeletal muscle health and NMJ genes in our entire clinical cohort (DM1 and Con; Table S6). Our analysis revealed that higher CARM1 activity was positively (*P < 0.05*) correlated with *CHRNA1, CHRND, CHRNE, MUSK* and *NCAM1* (Figure 6F). More importantly, ADMA^CARM1^ was negatively (*P < 0.05*) associated with skeletal muscle strength (i.e., pinch grip, knee extension, and grip strength), mass (i.e., appendicular and total lean mass), and fitness (i.e., VO_2_max, 6-minute walk test, forced expiratory volume, and forced vital capacity; Figure 6F).

**Figure 6.**
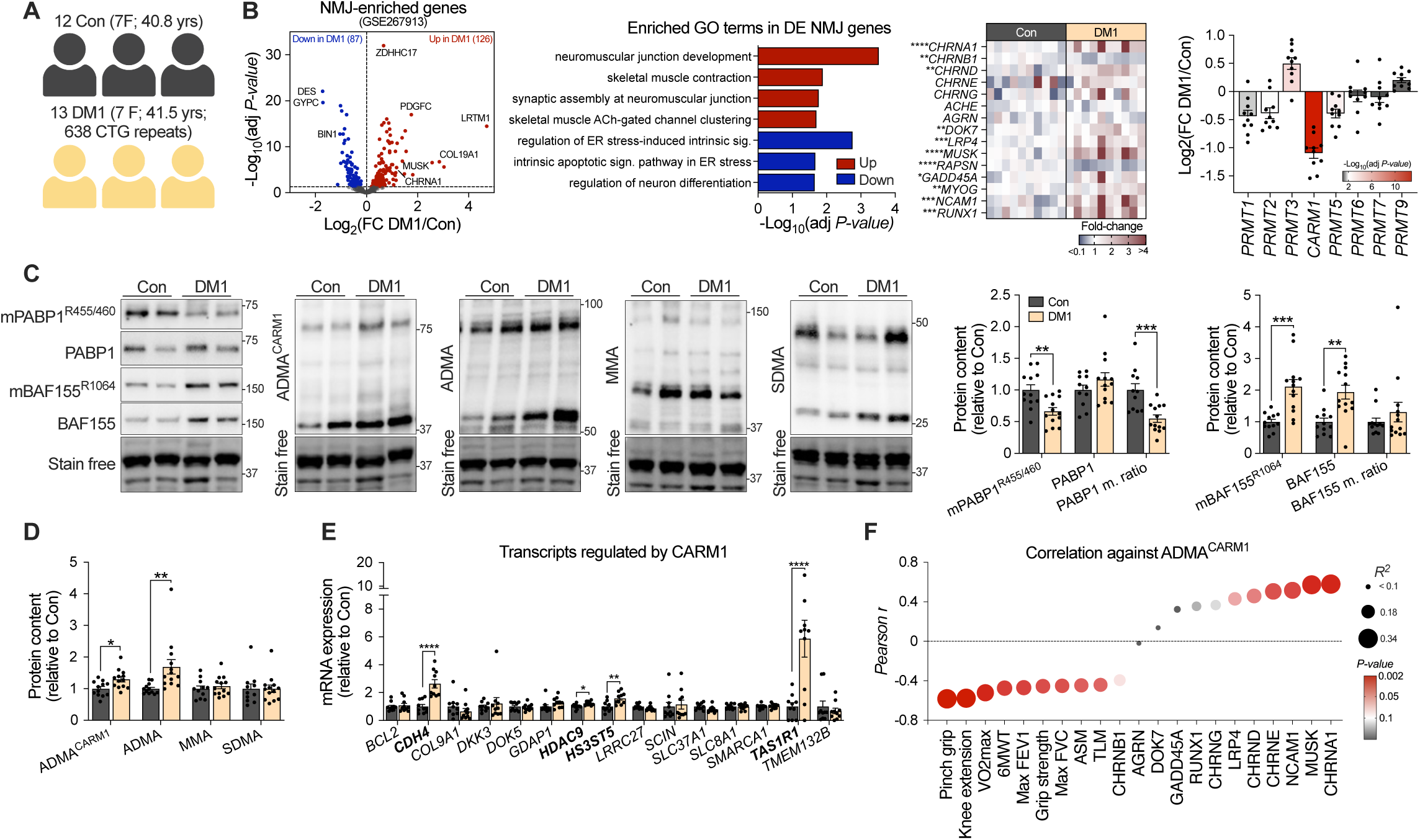
Increased global CARM1 activity in myotonic dystrophy type 1 (DM1) patients is associated with poor skeletal muscle mass and strength, and upregulated expression of NMJ transcripts. **(A)** Skeletal muscle biopsies from the vastus lateralis were sampled from 13 genetically confirmed myotonic dystrophy type 1 (DM1) patients along with 12 sex-, age- and body mass index (BMI)-matched unaffected controls (Con). **(B; Left)** Volcano plot indicating significantly upregulated (red) and downregulated (blue) NMJ-enriched transcripts in DM1 relative to Con. n = 10. **(B; Middle left)** GO pathway analysis performed on differentially expressed NMJ-specific genes. **(B; Middle right)** Heatmap summarizing mRNA expression of AChR subunits, denervation markers and other molecules important for NMJ health in skeletal muscle of Con and DM1 participants. n = 10. **(B; Right)** Total abundance of all *PRMTs* in skeletal muscle expressed as Log_2_(FC) DM1 relative to Con. n = 10. **(C; Left)** Representative Western blots for mPABP1^R455/460^, PABP1, mBAF155^R1064^, BAF155, ADMA^CARM1^, ADMA, MMA, and SDMA. Stain free blots displayed below show sample loading and approximate molecular weights (kDa) at right. **(C & D)** Graphical Summaries of PABP1 (methylated, total and methylation ratio), BAF155 (methylated, total and methylation ratio) and global arginine methylation markers. n = 12 for Con and 13 for DM1. **(E)** Total mRNA expression of transcripts potentially co-regulated in DM1 patient muscle by CARM1 that were acquired from bulk RNA sequencing of WT and CARM1 mKO skeletal muscle samples. **(F)** Bubble plot summarizing variables from a *Pearson’s* correlation analysis of ADMA^CARM1^ against pinch grip, knee extension, cardiorespiratory fitness (VO_2_max), 6-minute walk test (6MWT), forced expiratory volume (max FEV1), grip strength, forced vital capacity (max FVC), appendicular skeletal muscle mass (ASM), total lean mass (TLM), and transcripts encoding for AChR subunits (CHRNA1, CHRNB1, CHRND, CHRNE, and CHRNG), denervation markers (GADD45A, RUNX1, and NCAM1) and key NMJ regulators (DOK7, LRP4, and MUSK). Data are means ± SEM with individual data points displayed. \**P < 0.05*, \*\**P < 0.01*, \*\*\**P < 0.001*, *\*\*\*\*P < 0.0001* between groups. **(B & E)** A Benjamin-Hochberg (BH) multiple comparisons correction was applied and genes with an adj. *P-value* < 0.05 were considered significant. **(C & D)** A two-tailed unpaired Student T-test was used to calculate significance.

## DISCUSSION

Post-translational modification via arginine methylation has been generally understudied in skeletal muscle and neuromuscular biology as compared to other canonical protein modifications such as phosphorylation. Consequently, we and others have focused on uncovering novel roles for PRMTs within the peripheral neuromuscular system and found that they are required for optimal skeletal muscle size and function [9–13,33]. Here, we expanded on previous work and demonstrated that *Carm1* expression and methylation indicators of its activity are upregulated in NMJ-rich regions of mouse skeletal muscle, implicating methylarginine signalling in the maintenance of the NMJ. *Carm1* mKO led to larger, more fragmented NMJs and altered transcription of NMJ-specific genes under both basal and denervated conditions. Furthermore, we show that CARM1-mediated arginine methylation and transcription of a downstream target are increased in pre-clinical mouse models of DMD and SMA with characteristic NMJ instability, as well as during conditions of neuromuscular disuse and disease in humans, consistent with recent findings in ALS studies [18]. Finally, our data reveal that whole muscle CARM1 hyperactivity was associated with a more overt clinical phenotype and elevated expression of NMJ transcripts in DM1 patients. Collectively, these results provide strong evidence for CARM1 signalling in regulating NMJ stability, architecture and transcription during health and disease.

Maintenance and remodelling of the NMJ occurs, in part, via well-described mechanisms consisting of a series of tyrosine phosphorylation events in the Agrin-MuSK-Dok7 signalling axis to drive subsynaptic AChR expression and clustering [2,3]. However, numerous aspects of NMJ biology remain largely undefined. For example, recent evidence revealed that MuSK agonists elicit hundreds of previously unexplored phosphoproteomic signals in cultured muscle, which highlights our limited understanding of the complex mechanisms that govern the neuromuscular synapse [34]. In the present study, we provide evidence for the accumulation of CARM1-specific substrates and methylarginine residues at the motor endplate, which suggests a novel role for arginine methylation in NMJ biology. Previous work from our laboratory defined the arginine methylome in mouse skeletal muscle tissue [11]. While the majority of detected proteins were related to sarcomeric components and RNA-binding processes, we identified peptides localized to the NMJ, including LIM domain-binding 3 (Lbd3) [S7], and others that have been implicated in NMJ dysfunction such as fused in sarcoma (FUS) [S8] and Ataxin-2 [S9]. Nevertheless, the highly specialized organization and the low abundance of NMJ machinery present significant limitations for identifying key arginine methylation sites that may be important for the NMJ. Thus, future work should investigate the CARM1-dependent and -independent arginine methylproteome in NMJ-enriched samples to further expand our understanding of the mechanisms that regulate its development and remodelling.

We and others have shown that CARM1 is central to the regulation of skeletal muscle mass and function [9–12]. In the present study, we found that muscle-specific depletion of *Carm1* induces an adult-to-fetal shift in expression of AChR subunits (i.e., decreased *Chrnε* and increased *Chrnγ*) in NMJ-enriched regions and disrupts NMJ integrity, which, in part, could contribute to the atrophic muscle phenotype observed in these animals. Despite well-characterized phenotypic consequences of CARM1 deletion, the direct mechanisms by which CARM1 affects NMJ architecture and function remain unclear. Other members of the arginine methyltransferase family, such as PRMT1, are crucial for NMJ recovery following injury via a mechanism involving mitochondrial turnover [13,14]. Similarly, we have demonstrated that *Carm1* null skeletal muscle accumulates morphologically distorted mitochondria, and its expression is required for normal rates of mitophagic flux [12,25], providing a potential explanation for the NMJ dyshomeostasis in mKO mice. It’s important to note, however, that a skeletal muscle phenotype induced by blunted arginine methylation occurs following long-term germline deletions of PRMTs but not after short-term pharmacological inhibition during adulthood [9,10,33]. Therefore, time course investigations from development through early adulthood, as well as the use of inducible *Carm1* KO models, prior to onset of atrophy are necessary to decipher the complex interaction between pathways responsible for mitochondrial biology, the NMJ and muscle mass maintenance.

An alternative mechanism for CARM1-mediated effects on the NMJ involves its function as a transcriptional co-activator thereby modulating the expression of transcripts localized to the synapse. This hypothesis is reflected through the genotype-specific differences seen in NMJ genes following 7 days of denervation. We identified *Lrtm1* amongst the transcripts that were only upregulated in denervated WT animals, which was recently shown to augment muscle mass and exercise performance when depleted at a whole-body level but played a negligible role following its manipulation in skeletal muscle [26,35]. In contrast, *T-box transcription factor 21* (*Tbx21*) was preferentially downregulated in mKO-Den animals. Expression of *Tbx21* is required for NMJ recovery and reinnervation to restore muscle contractility after nerve damage, in part through the recruitment of immune cells [36]. Other molecules influenced by *Carm1* ablation include *disrupted in schizophrenia 1* (*Disc1*) and *cyclin-dependent kinase 6* (*Cdk6*), which are involved the Wnt/β-catenin and Notch signalling pathways, respectively, in non-muscle cells [S10–S12]. Both Wnt/β-catenin and Notch play important roles in the formation and plasticity of the NMJ [1,37–39]. Therefore, our transcriptomic analysis provides a preliminary framework for understanding CARM1’s influence on transcription of the NMJ machinery. Additional studies are required to elucidate the direct functional relevance of CARM1 at the NMJ.

Skeletal muscle displays robust increases in methylarginine signalling during neuromuscular remodelling elicited by synergistic ablation and neurogenic-induced atrophy [9,10,18]. Similar observations were made in ALS patients when assessing the whole muscle proteome [18]. Our results expand on the clinical significance of these previous studies and support the idea that enhanced CARM1-mediated arginine methylation is a shared mechanism across NMDs with varying genetic origins in mouse and human skeletal muscle. The hyperactivity of CARM1 in SMA and D2.*mdx* mice, which present with distinct but prominent NMJ defects, along with analysis of neuromuscular disuse in healthy humans, affirms a close relationship between the onset of NMJ instability and increases in CARM1 signalling. Moreover, inverse relationships between ADMA^CARM1^ content and metrics of skeletal muscle health, as well as positive correlations with NMJ molecules, present an intriguing hypothesis that targeted inhibition of the enzyme may provide therapeutic value for NMDs, potentially through NMJ-specific mechanisms. On the other hand, deleterious outcomes observed with complete suppression of CARM1 activity in skeletal muscle imply that stimulation of the methyltransferase may be protective during conditions of neuromuscular dysfunction. Future work should aim to reestablish hormetic signalling by partially inhibiting CARM1 in pre-clinical models of NMDs and other NMJ disorders to determine whether the associated upregulation of arginine methylation is adaptive or maladaptive. Additionally, the disconnect between total CARM1 protein and changes in its methylarginine transferase activity suggests an influence of upstream co-factors and requires further investigation.

In conclusion, we executed a series of complementary experiments aimed to address the role of CARM1 signalling at the neuromuscular synapse during health and disease. We highlight that CARM1 arginine methylation is preferentially enhanced in NMJ-enriched regions and that skeletal muscle-specific ablation of the enzyme elicits NMJ dysmorphia partially through dysregulated transcription of key synaptic mRNAs. Furthermore, the data reveal that changes in the expression of NMJ-specific genes after denervation are, in part, dependent on post-synaptic CARM1 activity. In skeletal muscle from mice and humans with the most prevalent NMDs, we established a mechanistic interaction between NMJ instability and CARM1-dependent arginine methylation, which persisted under non-pathological conditions in immobilized young healthy volunteers. Finally, we identified novel associations between CARM1-mediated methylarginine activity and clinically relevant measures of skeletal muscle health in adults with the most common form of muscular dystrophy. This study establishes that CARM1 is a key regulator of NMJ biology in health and disease and that targeting the enzyme may be beneficial in conditions of neuromuscular dysfunction.

## ACKNOWLEDGMENTS

We thank Dr. Mark Bedford (MD Anderson Cancer Center, University of Texas) for the *Carm1* floxed mice and ADMA^CARM1^ antibody. We are grateful to the technicians at McMaster University’s Central Animal Facility for routine animal husbandry, Todd Prior for his technical assistance and members of Exercise Metabolism Research Group for their support in this project. This work was supported by the Natural Science and Engineering Research Council of Canada (NSERC), Canadian Institutes of Health Research, Muscular Dystrophy Canada, Canada Research Chairs program, and the Ontario Ministry of Economic Development, Job Creation and Trade (MEDJCT). AIM and RR were recipients of an Ontario Graduate Scholarships. SYN, SRM, and DWS held NSERC Postgraduate Scholarships.

