## Supplemental information for "Arginine methyltransferase signalling is hyperactive in conditions of neuromuscular junction instability and muscle atrophy"

### SUPPLEMENTARY FIGURES

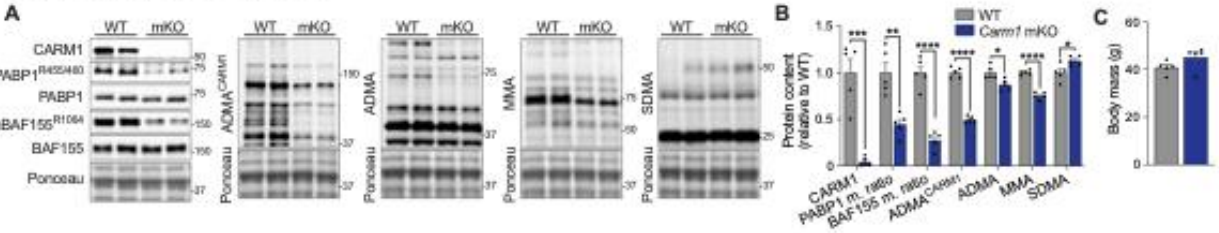


**Supplementary Figure 1. CARM1 mKO reduced downstream methylation targets but did not impact body mass. (A)** Representative Western blots for CARM1, mPABP1$^{R455/460}$, PABP1, mBAF155$^{R1064}$, BAF155, ADMA$^{CARM1}$, ADMA, MMA, and SDMA in TA muscles, along with **(B)** a corresponding graphical summary. Ponceau S stains displayed below show sample loading and approximate molecular weights (kDa) are at right of blots. n = 5. **(C)** Body mass of WT and mKO mice. n = 5. Data are means ± SEM with individual data points displayed. **P < 0.05*, ***P < 0.01*, ****P < 0.001*, *****P < 0.0001* between groups. Two-tailed unpaired Student T-test was used to calculate significance.

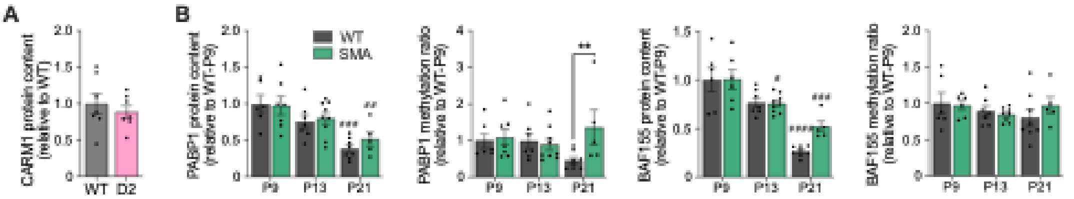


**Supplementary Figure 2. Corresponding quantification of CARM1 protein and downstream targets in pre-clinical models of NMDs. (A)** Quantification of total CARM1 protein expression in WT and D2.*mdx* TRI muscles relating to Figure 4B. n = 5-9. **(B)** Quantification of total and methylation ratio of PABP1 and BAF155 protein content in WT and SMA TRI muscles relating to Figure 4E & 4F. n = 5-9. Data are means ± SEM with individual data points displayed. ***P < 0.01* between different genotypes. $^{\#}$*P < 0.05*, $^{\#\#}$*P < 0.01*, $^{\#\#\#}$*P < 0.001*, $^{\#\#\#\#}$*P < 0.0001* relative to P9 within the same genotype. **(A)** A two-tailed unpaired Student T-test and **(B)** a two-way ANOVA followed by a Tukey or Šídák's post hoc test, when appropriate, were used to calculate significance.

### METHODS

*Animals. Carm1* fl/fl mice were recovered from cryopreserved sperm (a kind gift from Dr Mark Bedford; The University of Texas MD Anderson Cancer Center) and bred against transgenic animals expressing Cre recombinase under the control of the HSA promoter (Strain #: 006149, Jackson Laboratory) to generate *Carm1* mKO mice. Genotypes were confirmed by the presence of the *Carm1* fl/fl and Cre sequences in DNA isolated from ear notches using primers listed in Table S1. [1,2] DBA/2J-*mdx* (D2.*mdx*; Strain #: 013141) and WT DBA/2J (Strain #: 000671) mice were obtained from Jackson Laboratory and bred in-house as we have previously done [3,4]. To generate SMA mice and their WT counterparts, $Smn^{2B/2B}$ animals were crossed with heterozygous $Smn^{+/-}$ mice to generate healthy $Smn^{2B/+}$ (WT) and $Smn^{2B/-}$ (SMA) mice, which mimic a Type II SMA phenotype [5–7]. We used both male and female mice for SMA experiments as they present similarly in phenotype and disease progression [8].

*Single leg immobilization experiment.* Briefly, 24 young healthy men (22 ± 2 yrs old) with a BMI ranging from 20-30 kg/m$^2$ were recruited to McMaster University as part of a previous investigation [9]. Following initial health screening, participants underwent a baseline skeletal muscle biopsy (Pre) from the vastus lateralis after which they were subjected to 14 days of single leg immobilization using a fixed knee brace. A final muscle biopsy (Post) sample was collected after the immobilization period. Both Pre and Post biopsy samples from the same participant were available for 18/24 and 15/24 subjects to perform mRNA and protein analysis, respectively.

*DM1 and unaffected controls.* A total of 13 genetically confirmed DM1 patients (41.5 ± 3 yrs old) and 12 matched healthy controls (Con; 40.8 ± 3 yrs old) were recruited from the Neuromuscular and Neurometabolic Clinic at McMaster University. Resting skeletal muscle biopsies from the vastus lateralis were obtained using a modified Bergström needle [10]. All clinical metrics of strength, muscle mass and function used for *Pearson's* correlation analysis were completed as outlined previously [11]. RNA sequencing (GSE184951) was available for 10 DM1 and 10 Con participants, while protein samples were available for the entire cohort.

*Sciatic nerve transection.*Click or tap here to enter text.. Animals were subcutaneously injected with anafen (2 mg/kg) prior to being anaesthetized under 2-3% of isoflurane. A small skin incision was made, and the underlying muscles were gently separated with blunt forceps to expose the sciatic nerve. Approximately 0.5 cm of the nerve was excised, after which the incision was sutured, and animals were allowed to recover over a heating pad. The TA muscles were collected 7-days post transection from both the Den and contralateral Con limbs for RNA sequencing.

*RNA isolation, purification and quantitative real-time polymerase chain reaction (qPCR).* Muscle samples were homogenized in 1 mL of TRIzol reagent (Invitrogen). The RNA phase was purified using the E.Z.N.A RNA kit I (R6834-02, Omega Bio-Tek kit), samples were adjusted to a concentration of 200 ng/µL and reverse transcribed using a high-capacity cDNA reverse transcription kit (4368814, Thermo Fisher Scientific) as instructed by the manufacturer. All samples were run in triplicate on a 384-well plate with each reaction containing 2 μg of cDNA and GoTaq qPCR Master Mix (A6002, Promega). Gene expression was determined using the comparative $C_T$ method [13]. Expression of *ribosomal protein S11* (*Rps11;* mouse experiments) and *ribosomal RNA 18S* (human experiments) did not differ between groups and were therefore used as housekeeping genes. All qPCR primers used are listed in Table S1.

*Whole-mount immunofluorescence (IF) NMJ staining and analysis.* Morphology of the pre- and post-synapse was carried out as previously described [14]. Whole ETA muscles were dissected, pinned in a Sylgard-coated 10 mm Petri dish, fixed at room temperature (RT) with 4% PFA for 10 mins and stored in PBS at 4 °C for further processing. Muscles were then neutralized in 0.1M glycine, permeabilized with 0.5% Triton X-100 and incubated in blocking solution consisting of 3% bovine serum albumin (BSA), 5% goat serum and 0.5% Triton X-100 in 1X PBS. To identify the motor neuron axon and terminal, samples were incubated overnight at 4 °C with antibodies against NF and Syn, respectively (Table S2). Next day, samples were incubated in the appropriate secondary antibodies and AChRs were labeled with α-BTX Alexa Fluor (AF)-488 (1:500, B13422, Thermo Fisher Scientific). Finally, samples were allowed to dry fully and mounted with Prolong Gold antifade reagent (P36930, Thermo Fisher Scientific) before images were captured via a 60x Plan Apochromat 1.4 NA oil immersion objective (Nikon Instruments), Nikon C2$^{+}$ confocal microscopy unit (Nikon Instruments), and a Nikon LU-N4 laser system (Nikon Instruments). A minimum of 15 NMJs were analyzed per sample in a blinded fashion using NMJ-morph as previously described [15].

*Protein isolation and Western blotting.* Flash frozen muscle samples were placed in RIPA buffer (1:20, R0279, Sigma-Aldrich) supplemented with phosphatase inhibitors (4906837001, Roche) and protease (04693159001, Roche) and homogenized using a motorized tissue lyser

(Qiagen). Protein concentrations were assessed using a bicinchoninic protein assay (BCA; PI23225, Thermo Fisher Scientific). For Western blotting, proteins (~20 μg) were separated on a Criterion TGX precast gel (5671095, Bio-Rad Laboratories) and transferred onto nitrocellulose membranes (1620112, Bio-Rad Laboratories). Blots were blocked in 5% BSA in 1X TBST for 1 hour prior to an overnight incubation at 4 °C with primary antibodies listed in Table S2 and visualized using chemiluminescence reagent (1705061, Bio-Rad Laboratories). Images were quantified using Image Lab software (Bio-Rad Laboratories) and all bands were normalized to whole-lane protein content as determined by ponceau staining or stain-free intensity.

*RNA sequencing and analysis.* RNA sequencing was carried out as previously described by our group [16,17]. Total RNA was extracted as described in the aforementioned section and only samples with a RNA quality number (RQN) of 8.0 or greater were used for library construction. Lowly expressed transcripts (<10 counts across all samples) were filtered and differential expression analysis was executed using DESeq2. A Benjamin-Hochberg (BH) correction was applied to account for multiple comparisons and only genes with an adjusted *P-value* < 0.05 were considered significant for subsequenty analysis. Gene ontology (GO) pathway enrichment analysis was performed on significant transcripts using clusterProfiler. A list of NMJ-specific transcripts previously identified by Ham et al [18] using sn-RNA sequencing was used as a reference in the present study. All RNA sequencing analyses were performed using R Studio [9,11].
